# Extreme positive allometry of animal adhesive pads and the size limits of adhesion-based climbing

**DOI:** 10.1101/033845

**Authors:** David Labonte, Christofer J. Clemente, Alex Dittrich, Chi-Yun Kuo, Alfred J. Crosby, Duncan J. Irschick, Walter Federle

## Abstract

Organismal functions are size-dependent whenever body surfaces supply body volumes. Larger organisms can develop strongly folded internal surfaces for enhanced diffusion, but in many cases areas cannot be folded so that their enlargement is constrained by anatomy, presenting a problem for larger animals. Here, we study the allometry of adhesive pad area in 225 climbing animal species, covering more than seven orders of magnitude in weight. Across all taxa, adhesive pad area showed extreme positive allometry and scaled with weight, implying a 200-fold increase of relative pad area from mites to geckos. However, allometric scaling coefficients for pad area systematically decreased with taxonomic level, and were close to isometry when evolutionary history was accounted for, indicating that the substantial anatomical changes required to achieve increases in relative pad area are limited by phylogenetic constraints. Using a comparative phylogenetic approach, we found that the departure from isometry is almost exclusively caused by large differences in size-corrected pad area between arthropods and vertebrates. To mitigate the expected decrease of weight-specific adhesion within closely related taxa where pad area scaled close to isometry, data for several taxa suggest that the pads’ adhesive strength increased for larger animals. The combination of adjustments in relative pad area for distantly related taxa and changes in adhesive strength for closely related groups helps explain how climbing with adhesive pads has evolved in animals varying over seven orders of magnitude in body weight. Our results illustrate the size limits of adhesion-based climbing, with profound implications for large-scale bio-inspired adhesives.

---

The evolution of adaptive traits is driven by selective pressures, but can be bound by phylogenetic, developmental and physical constraints [1]. Integrating evolution and biomechanics provides a powerful tool to unravel this complex interaction, as physical constraints can often be predicted easily from first principles [2]. The influence of physical constraints is especially evident in comparative studies across organisms which differ substantially in size [3,4,5,6]. For example, Fick’s laws of diffusion state that diffusive transport becomes increasingly insufficient over large distances, explaining the development of enlarged surfaces for gas and nutrient exchange (e. g. leaves, roots, lungs, gills, guts) and integrated long-distance fluid transport systems (e. g. xylem/phloem, circulatory systems) in larger animals and plants. How these systems change with size is determined by physical constraints [7,8,9]. While ‘fractal’ surface enlargements are possible without disrupting other body functions, strong positive allometry can conflict with anatomical constraints. For example, structural stability demands that animals should increase the cross-sectional area of their bones in proportion to their body weight, but excessively thick leg bones can compromise other physiological functions and hamper locomotion [3,10,11].

Adhesive pads are another example of an adaptive trait subject to size-dependent physical constraints. These systems allow animals to climb smooth vertical or inverted surfaces, thereby opening up new habitats. Adhesive pads have evolved multiple times independently within arthropods, reptiles, amphibians and mammals, and show impressive performance: they are rapidly controllable, can be used repeatedly without any loss of performance, and function on rough, dirty and flooded surfaces [12]. This performance has inspired a considerable amount of work on technical adhesives that mimic these properties [13]. A key challenge for both biological and bio-inspired adhesive systems is to achieve size-independent performance [14, 15, 16], i.e. the maximum sustainable adhesion force, *F*, should be proportional to the mass to be supported, m. For vertically climbing animals, *F* is the product of the maximum adhesive stress, *σ*, and the adhesive pad area, *A*, each of which may change with mass *(A* oc *m^a^* and *σ* ∝ *m^b^*), so that constant size-specific attachment performance requires:

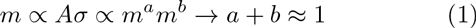

where *a* and *b* are the scaling coefficients for *σ* and *A* in relation to body mass, respectively. If animals maintain geometric similarity when increasing in size, *A* would scale as m^2^/^3^, so that the adhesion per body weight for large geckos (m ≈100 g) is expected to be approximately 10^7^/^3^ ≈200 times smaller than for tiny mites (m ≈10*µ*g) if the pads’ adhesive strength *σ* remained unchanged (b = 0). Large animals can only circumvent this problem by (i) developing disproportionally large adhesive pads (a > 2/3), and/or (ii) systematically increasing the maximum force per unit pad area (b > 0). How do large climbing animals achieve adhesive forces equivalent to their body weight?

Using the simple biomechanics argument outlined above as a framework, we here provide a comparative analysis of the allometry of adhesive pad area across 225 species, covering more than seven orders of magnitude in weight – almost the entire weight range of animals climbing with adhesive pads — and including representatives from all major groups of adhesion-based climbers.

**Figure 1.**
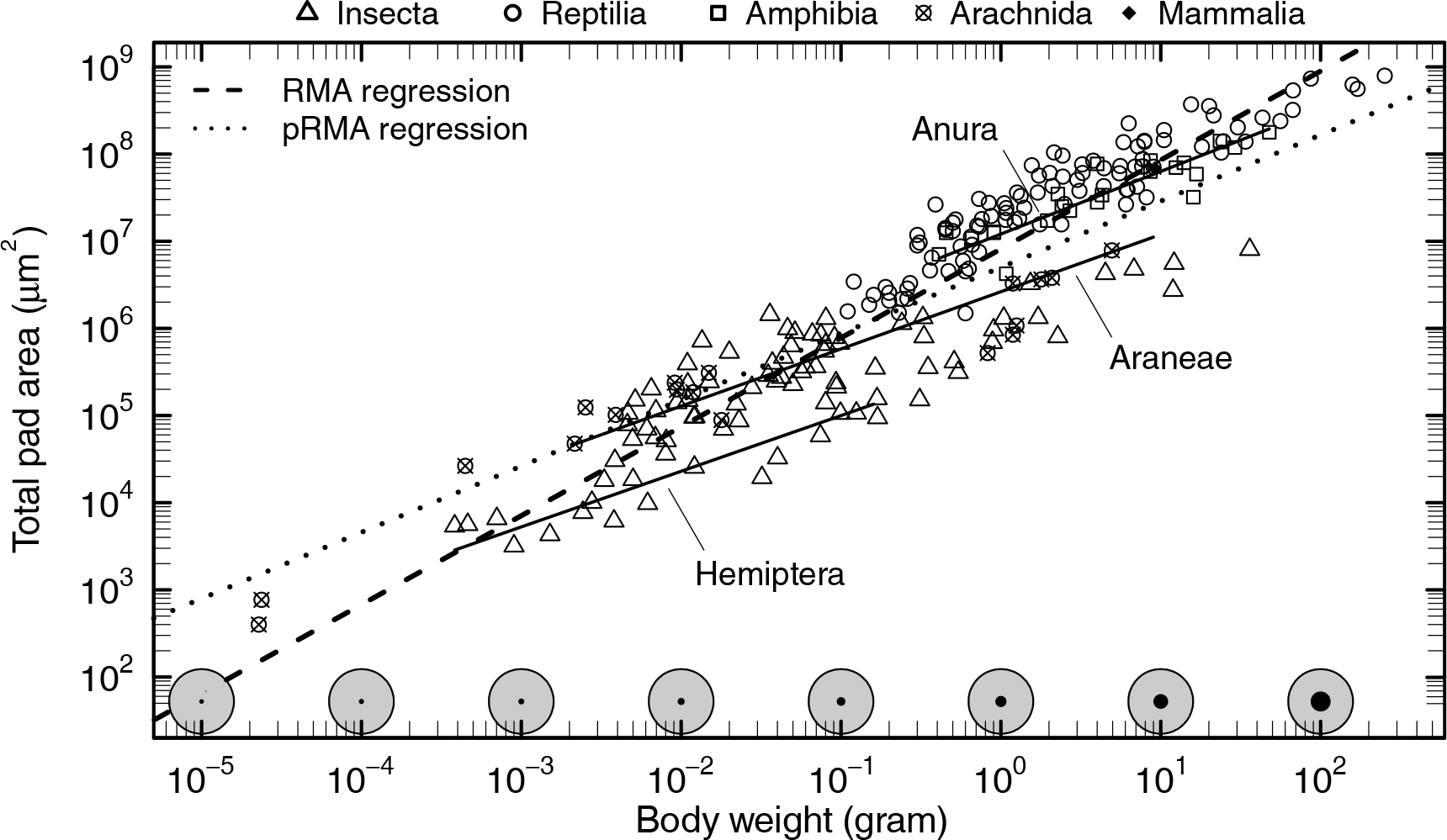
Allometry of pad area for 225 species, ranging from ≈20*µ*g to ≈200g in body weight. Across all taxa, pad area was directly proportional to body weight (reduced major axis regression, RMA, dashed line). The increase in the fraction (black circles) of the total available surface area (grey circles) required to accommodate this disproportionate change is schematically illustrated along the bottom axis, assuming that a climbing animal of 80kg has a total body surface area of 2m^2^, comparable to a human of 80kg and 180 cm, and that body surface area is approximately isometric. Strikingly, scaling relationships were significantly less steep within closely related groups (representative solid lines from RMA regressions), and closer to isometry when phylogenetic relationships were accounted for (phylogenetic RMA regression, dotted line).

## Results and Discussion

### Scaling of adhesive pad area

Across all taxa, adhesive pad area showed extreme positive allometry and scaled as A ∝ m^1.02^ (reduced major axis regression (RMA); see fig. 1 and tab. 1 for detailed statistics), an increase sufficient to compensate for the predicted loss of weight-specific adhesion, even if adhesive strength remained unchanged. Thus, adhesive pads occupy a larger fraction of the body surface area in larger animals. Within closely related taxonomic groups, however, pad area grew more slowly with body mass, indicating a strong phylogenetic signal (see fig. 1 and 2).

When evolutionary relationships were accounted for, the observed scaling coefficient decreased dramatically, and was consistent with isometry (fig. 1, 2 and tab. 1). This systematic change of allometric coefficients with taxonomic rank suggests that phylogenetic inertia impedes a disproportionate increase of pad area within closely related groups (fig.2 and 3A). Our results thus add to a body of evidence suggesting that the evolutionary flexibility of allometric slopes is low and larger changes in particular traits are mainly achieved by shifts of the allometric elevation [18,19].

Removal of the influence of body size by analysing the residuals (termed ‘relative pad area’ in the following) of a phylogenetic reduced major axis regression (pRMA) allowed us to further investigate at what taxonomic level major shifts in relative pad area occurred, separating the effects of size and ancestry. Relative pad area differed strongly between vertebrates and arthropods, but comparatively little variation existed within these groups (fig. 3A). More than 58% of the variation in residual pad area was explained by differences between vertebrates and arthropods (nested ANOVA, F_1,173_=845, p<0.001, see tab. 2), so that body weight and phylum alone accounted for more than 90% of the total variation in pad area. Rather than being driven solely by variation in body size, differences in relative pad area appear to be tied to characteristic features of the corresponding phyla, such as, for example, the presence or absence of multiple distal pads (toes) per leg (fig.3B,C). However, we also found evidence for differences in relative pad area within lower taxonomic ranks. For example, members of the gecko genus *Sphaerodactylus* had considerably smaller pads than other Gekkotan lizards, whereas *Gekko* lizards had particularly well-developed pads (based on their relative pad area, fig.3 A). In insects, hemimetabolous orders had smaller relative pad areas than holometabolous orders (see fig.3A). Adhesive pads can allow access to arboreal habitats [20, 21, 22], but they may come at the cost of reduced locomotor performance in situations where no adhesion is required [23,24]. Thus, the multiple independent losses, gains, and reductions of adhesive pads in amphibians, insects, lizards and spiders [25, 26, 27, 28] likely reflect the ecological, behavioural and taxonomic diversity within these groups [29,30].

**Table 1.** Results for generalised least squares and reduced major axis regressions describing the relationship between log_10_(adhesive pad area) (in *μm^2^*) and log_10_(mass) (in g) across all taxa. Covariance in pad area and body mass between related species was either ignored (uncorrected) or accounted for (corrected). Pagel’s lambda is a statistic measuring the strength of phylogenetic signal (λ=1 indicates that the trait evolves like Brownian motion along the phylogeny, whereas λ=0 indicates that the trait is not correlated with phylogeny [17].). Numbers in brackets give approximate 95% confidence intervals of the estimated parameters where available.

| Uncorrected | Elevation | Slope | Pagel’s $\lambda$ |
| --- | --- | --- | --- |
| Reduced major axis |  |  |  |
| Pad area against mass | 6.91 (6.84, 6.98) | 1.02 (0.97, 1.07) | 0 (fixed) |
| Mass against pad area | -6.76 (-7.08, -6.44) | 0.98 (0.93, 1.03) | 0 (fixed) |
| Generalised least squares |  |  |  |
| Pad area against mass | 6.88 (6.80, 6.94) | 0.95 (0.9, 1) | 0 (fixed) |
| Mass against pad area | -6.29 (-6.61, -5.96) | 0.91 (0.86, 0.95) | 0 (fixed) |
| Corrected | Elevation | Slope | Pagel’s $\lambda$ |
| Reduced major axis |  |  |  |
| Pad area against mass | 6.54 (6.47, 6.62) | 0.78 (0.74, 0.83) | 0.93 (fitted) |
| Mass against pad area | -8.38 (-8.77, -7.99) | 1.28 (1.21, 1.36) | 0.93 (fitted) |
| Generalised least squares |  |  |  |
| Pad area against mass | 6.44 (6.28, 6.61) | 0.70 (0.66, 0.75) | 0.90 (fitted) |
| Mass against pad area | -7.54 (-7.97, -7.10) | 1.13 (1.06, 1.20) | 0.83 (fitted) |

### The size-limits of adhesion-based climbing

Strong positive allometry of non-convoluted body structures in organisms ranging in size over many orders of magnitude is difficult to achieve, owing to simple anatomical constraints. For example, bone mass in terrestrial animals is predicted to increase with mass^4/3^ to maintain constant bone stress levels, but this would require unrealistic relative bone masses for larger mammals (scaling up an 8-gram shrew with ca 4% bone mass would produce a rather unfortunate 8-tonne elephant with 400% bone mass). The actual scaling coefficient is inevitably smaller (≈ 1.1) [10, 31], and alternative strategies have evolved to limit bone stresses [11].

Maintaining a pad area proportional to body weight in animals ranging from 10^−5^ to 10^2^ grams requires extraordinary morphological changes: assuming otherwise isometric animals, the proportion of the total body surface area specialised as adhesive pad needs to increase by a factor of 10^7/3^ ≈ 200. This extreme shape change may impose a size limit for adhesion-based climbing. Scaling up the relative pad area of arthropods and small vertebrates to a human of 180cm body length and 80kg body mass would result in an adhesive pad area of ≈ 10^6.91^·80000^1.02^ ≈ 0.81 m^2^, approximately 2/5 of the total available body surface area (≈ 2 m^2^, [33]). The required morphological changes, if at all possible, would thus be enormous, and difficult to achieve over short evolutionary timescales. Our results therefore indicate that phylogenetic inertia restricts the ‘design space’ for evolution at least for closely related taxa. Larger animals within closely related taxa must therefore either cope with a size-related decrease in their relative attachment ability, or develop alternative strategies to compensate for it. Recent studies on tree frogs and ants revealed that pad adhesive strength can vary systematically with size, resulting in an almost body size-independent attachment abilities despite a near-isometric growth of pad area [16,34]. Here, we extend these studies to investigate whether such adaptations are also present above species or genus level.

Figure 4 shows whole-body adhesion per pad area plotted against body weight for 17 frog species from 4 families and 12 genera [35,36,37,38]. All adhesion measurements were conducted using a tilting platform, and are thus comparable across studies. Over two orders of magnitude in body weight, adhesion force per unit area increased with m^0.3^ (RMA slope 0.3, 95% confidence interval (CI): (0.2, 0.43), generalised least-squares (GLS) slope 0.19, 95% CI: (0.08, 0.31)), sufficient to achieve body size-independent adhesive performance despite an approximately isometric growth of pad area (RMA slope 0.74, 95% CI: (0.62, 0.87), GLS slope 0.70, 95% CI: (0.58, 0.83)). In contrast to our results for the allometry of pad area, this relationship remained virtually unchanged when phylogenetic relationships were accounted for, indicating that pad performance directly responded to selective pressures unconstrained by phylogenetic history (phylogenetic RMA slope 0.28, 95% CI: (0.2, 0.42), phylogenetic GLS slope 0.17, 95% CI: (0.07, 0.27), see fig. 4). Together, our results provide strong evidence that two different strategies have evolved to deal with the physical challenges of a larger body size. These strategies have been adopted at different taxonomic levels, highlighting how phylogenetic and physical constraints can influence the evolution of adaptive traits. Across distantly related groups, leg morphology is sufficiently different to accommodate large differences in relative pad area. Within closely related groups, where anatomical constraints result in a scaling of pad area closer to isometry, some taxa appear to have increased their pads’ adhesive efficiency with size. The mechanisms underlying this increase in adhesive strength are still unclear, but may be of considerable interest for the development of large-scale bio-inspired adhesives. Various hypotheses have been proposed [14,16,37]. but still remain to be tested.

### Force scaling and the evolution of ‘hairy’ adhesive pads

Arzt et al (2003) suggested that large animals with hairy adhesive pads have evolved higher hair densities to increase their pads’ adhesive strength, an idea derived from the assumption that adhesive forces scale with the width of individual hair tips [39]. Assuming *isometric growth* of the total pad area, Arzt et al. predicted that hair density would need to increase with m^2/3^ in order to achieve constant mass-specific adhesion, in agreement with the data presented (but see ref. [40] which showed that adhesive hair density increased with body mass only when species were treated as independent data points, but not when phylogeny was considered). However, our data show that total pad area is *directly proportional* to body mass across distantly related taxa, so that a constant hair density would suffice. In addition, there is no experimental evidence that the adhesive strength of animal adhesive pads increases with decreasing size of individual contacts [16,41]. Thus, it appears unlikely that ‘force scaling’ has played an important role in the evolution of fibrillar adhesive systems [42].

**Figure 2.**
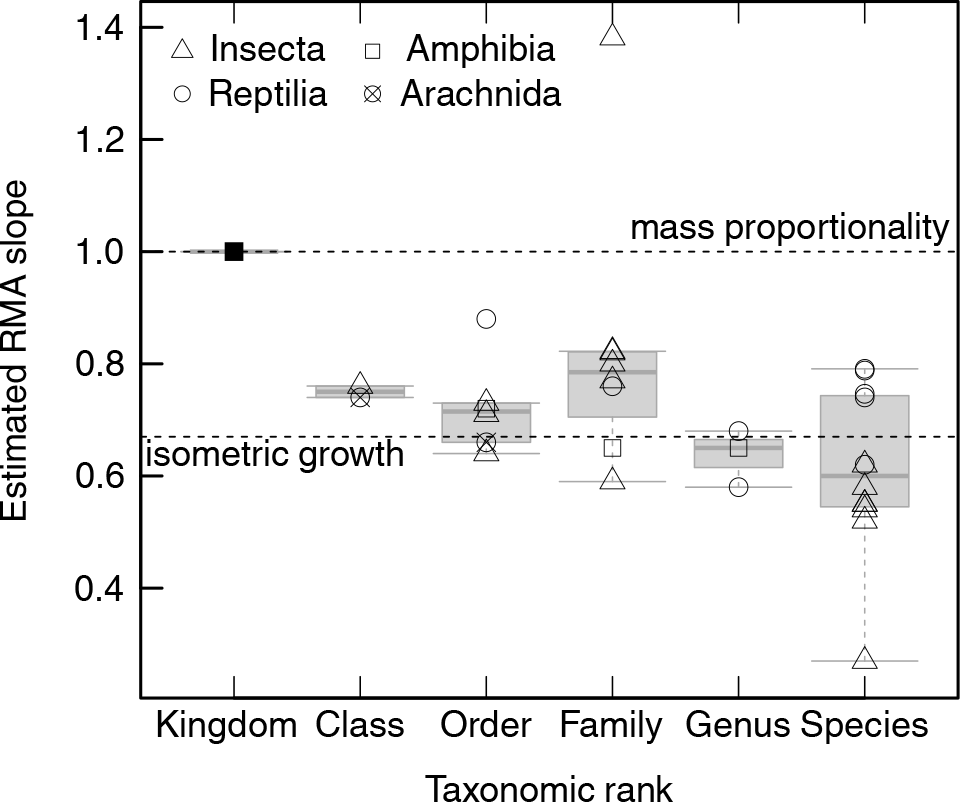
Change in pad allometry with taxonomic rank. For example, the slope of the regression line for Araneae in fig.1 is one data point for the rank ‘order’. Allometric coefficients decreased systematically, from mass-proportionality across all animals to isometry within genera and species. Data for the species-level allome-try are from [16,32].

Adhesive pads constitute a prime model system for studying the link between morphology, performance and fitness [43]. Further mechanistic and comparative studies are needed to elucidate the factors driving the evolution of these structures, and may ultimately allow us to mimic their properties with synthetic adhesives.

## Materials and Methods

### Data collection

Data were either collected by the authors or extracted from references [16, 29, 35, 36, 37, 38, 41, 44, 45, 46, 47, 48, 49, 50, 51, 52, 53, 54, 55, 56, 57, 58, 59, 60, 61, 62, 63, 64, 65, 66, 67, 68, 69, 70, 71, 72, 73, 74, 75, 76, 77, 78, 79, 80, 81, 82, 83, 84, 85, 86].

Arthropod specimens were collected around Cambridge (UK), Brisbane (Australia), or obtained from the Cambridge Phasmid Study Group. All arthropods were identified [87, 88, 89], and their live weight was recorded (ME5, resolution 1µg, max 5g or 1202 MP, resolution 0.01g, max 300g, both Sartorius AG, Goettingen, Germany). Attachment pads were photographed either with a Canon EOS (Canon, Tokyo, Japan) mounted on a stereo microscope (MZ16, Leica Microsystems Ltd., Heidelberg, Germany), or by using scanning electron microscopy (SEM) for large and small specimens, respectively. Some pads were imaged whilst in contact with glass, visualised using the stereo-microscope with coaxial illumination. For SEM imaging, individual legs were dried, mounted on stubs, sputter-coated at 65 mA for 10-20 s (K575X turbo-pump sputter, Quorum Technologies, Sussex, UK) and examined with a field emission gun SEM at a beam voltage of 5 kV (Leo Gemini 1530VP, Carl-Zeiss NTS GmbH, Oberkochen, Germany).

Data on toepad-bearing gecko species were collected from live animals kept in the D.J.I. laboratory (under an Institutional Animal Care and Use (IACUC) protocol 2012–0064 from the University of Massachusetts at Amherst to DJ Irschick), and preserved specimens from the American Museum of Natural History and the Museum of Comparative Zoology at Harvard University. For each specimen, photos of one fore foot were obtained by pressing it tightly against the glass plate of an Epson Perfection V500 Photo Scanner (Seiko Epson Corp., Owa, Suwa, Nagano, Japan) next to a ruler and taking a digital scan. The total toepad area across all digits was measured using ImageJ v1.49r [90]. We also measured snout-vent-length (SVL, ±1mm) from each individual using a clear plastic ruler. Where possible, we measured multiple conspecific individuals, and used the mean as the species value.

Literature data were taken from the papers’ text or tables, or were extracted from figures using Web-PlotDigitizer 3.3 (WPD, developed by Ankit Ro-hatgi (http://arohatgi.info/WebPlotDigitizer), or Im-ageJ v1.49m. We tested the performance of WPD with an x-y plot of 20 random numbers between 1 and 1000 (x) and 0.01 and 10 000 (y) on a log-log scale, and found an accuracy of ≈0.6% for the raw data.

We used live body weights where available, and interpolated live weight from body length where necessary, using established scaling relationships [40]. A list of the included species can be found in the appendix.

**Table 2.** Results for a nested ANOVA on the residuals of a phylogenetic reduced major axis regression. *η*^2^ is the variance in residual pad area accounted for the by the different taxonomic levels, and most of the variation occurs between phyla.

| Tax. level | df | Mean squares | F | p-value | $\eta^2$ |
| --- | --- | --- | --- | --- | --- |
| Phylum | 1 | 40.25 | 845.16 | < 0.001 | 0.58 |
| Class | 3 | 0.41 | 8.55 | < 0.001 | 0.02 |
| Order | 12 | 1.02 | 21.43 | < 0.001 | 0.18 |
| Family | 35 | 0.22 | 4.52 | < 0.001 | 0.11 |
| Residuals | 173 | 0.05 |  |  |  |

**Figure 3.**
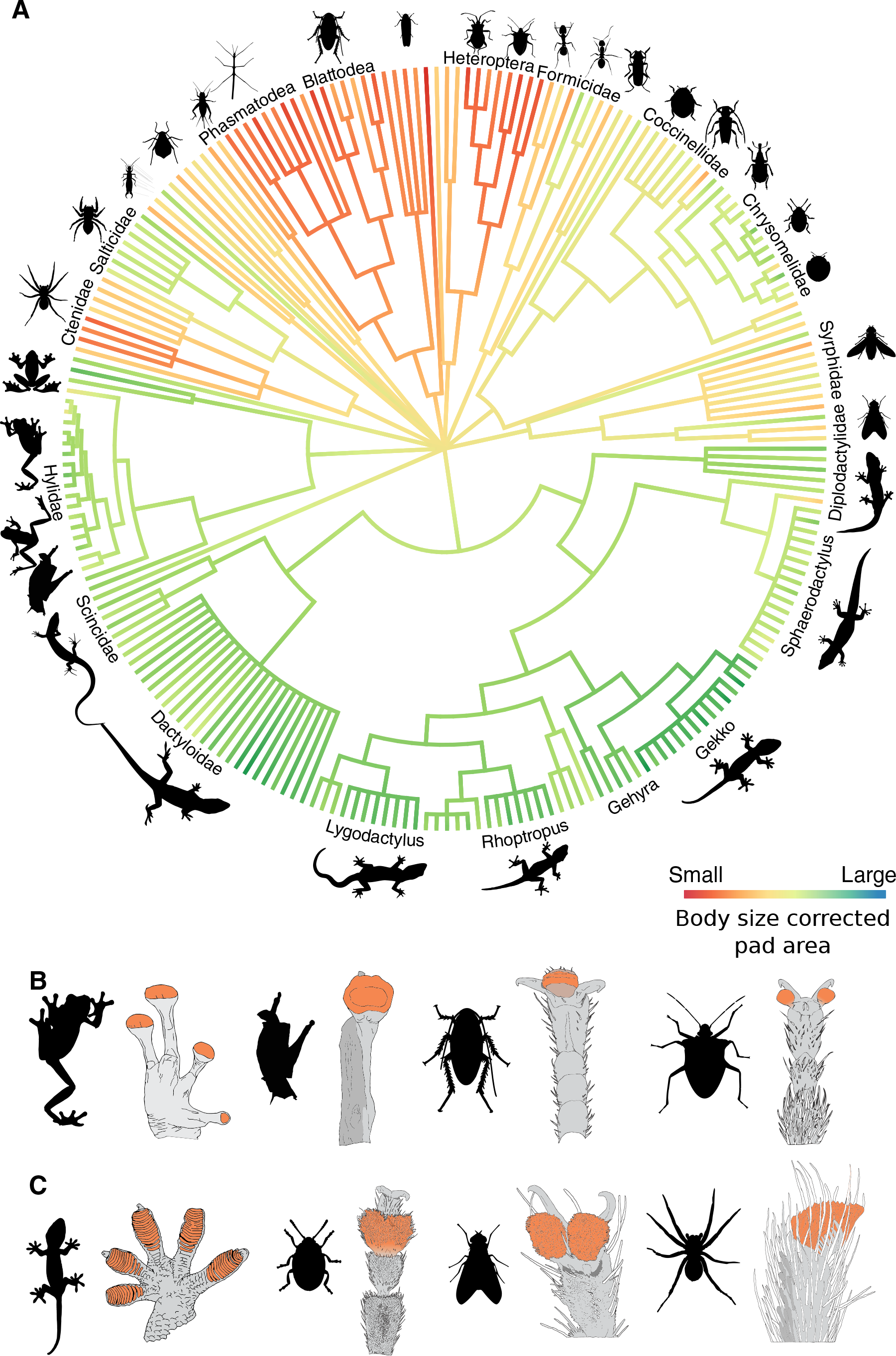
(A) Overview of the diversity of taxa examined in this study. Branch lengths do not reflect time or base pair substitutions (see methods). Branches are coloured according to a maximum likelihood estimate of the ancestral state of relative pad area (i. e. the residuals from a log-log regression of pad area against body weight), visualising systematic differences in relative pad area between arthropods and vertebrates. All values apart from tip states are only approximate and are not used to support any conclusions (see methods). (B & C) Cartoons depicting footpad morphology for representative groups within the phylogeny shown above with smooth and hairy adhesive pads. Projected pad area is highlighted in orange for each representative (see methods).

### Adhesive pad area

Animals attach to smooth surfaces by employing specialised attachment pads on their legs. These pads are either covered with dense arrays of fibrils (‘hairy pads’), or are macroscopically unstructured (‘smooth pads’). In order to compare pad areas across different taxa and pad morphologies, the following assumptions were made:

1. *‘Projected’ pad area is the most meaningful measure of contact area in comparative studies*. Projected pad area is the surface area of the foot specialised specifically for generating adhesion and friction [91]. In a fibrillar pad, inevitably only a fraction of this area comes into surface contact, i. e. the ‘real’ contact area is significantly smaller than the projected contact area.
2. *All animals employ a similar fraction of their available pad area when maximum performance is required*. It is unclear what fraction of the available pad area is employed by animals of different size [16, 92, 93], and systematic studies are lacking. However, the small number of direct contact area observations available strongly suggest that animals often use the entire area of their adhesive pads [91, 94], i. e. that they are not ‘overbuilt’. We thus assume that all climbing pads are designed so that their whole area can be used in critical situations.
3. *Adhesive performance is dominated by distal pads*. Insects can have several attachment pads per leg. There is strong evidence that these pad types differ in their morphology, as well as in their performance and function during locomotion [16, 44, 65, 70, 91]. Many insects do not employ their distal pads when no adhesion is required [44, 65, 95, 96, 97, 98], whereas during inverted climbing, only distal pads are in surface contact [83]. Accordingly, insects with ablated distal pads cannot cling upside down to smooth surfaces [44, 98]. Distal pads thus appear to be true ‘adhesive pads’ [83]. Proximal pads, in contrast, can be ‘non-sticky’, and may be designed as non-adhesive friction pads [99,100]. The distal pads are usually part of the pretarsus, but some insects lack a pre-tarsal pad. In these insects, the distal tarsal pad can show similar morphological specialisations [101,102]. Proximal pads are mainly found in arthropods, but they may also be present in frogs [35,103]. As the contribution of proximal pads to adhesion is unclear and likely variable, we exclude them for this study. In most insects, the total proximal pad area is between 3-5 times larger than the distal pad area, and pad area is still positively allometric even when proximal pads are included (reduced major axis regression slope between 0.8-0.9).
4. *The variation of pad area/adhesive strength between different legs/toes and sexes of the same species, and the variation introduced by the animals’ ecology is independent and randomly distributed with respect to body weight*. Several studies have shown that the size, morphology and performance of attachment devices can depend on the ecological niche occupied by the animals [20, 22, 29, 59, 104]. Variation can also occur between sexes [70, 105, 106], different legs or toes [20, 84], or even between populations of the same species occupying different habitats [24]. For this study, we assume that because of the large number of samples and the wide range of body sizes included, any bias introduced by these factors can be ignored.
5. *Adhesive performance of ‘wet’ and ‘dry’ pads is comparable*. Dynamic adhesive pads are frequently categorised as ‘wet’ or ‘dry’. However, there is no evidence for a functional difference between these two pad types [107], and indeed maximal adhesive stresses are comparable [16].

**Figure 4.**
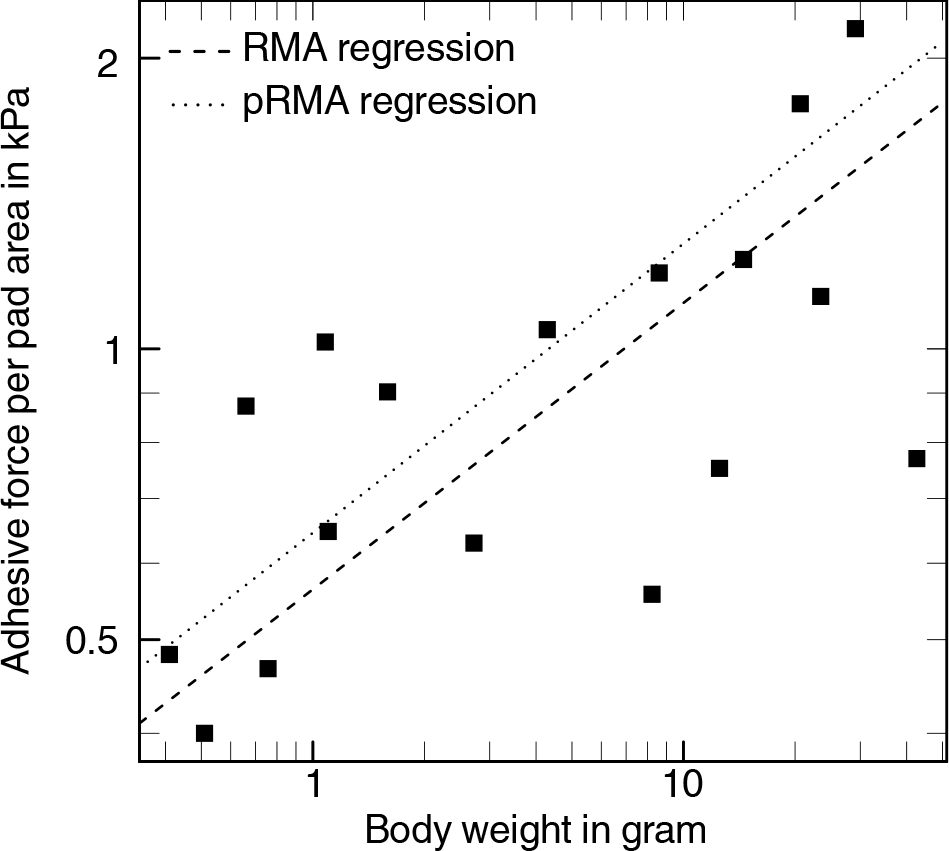
Adhesive force per area in 17 tree frog species increased with body mass, indicating that larger species possess more efficient pads. In contrast to the allometry of pad area, this result was not significantly influenced by phylogeny.

As some of the data used in this study originate from different groups, we quantified the consistency of pad area measurements among researchers. A selection of SEM images (three hairy, and seven smooth pads) were given to 10 scientists who independently measured nominal pad area. We found an average coefficient of variation of 17±9%, which was independent of the animals’ body weight (ANOVA, F_1,8_=0.067, p=0.8). Scaling relationships calculated with this dataset did not vary significantly across scientists (slope: likelihood ratio statistic= 0.38; elevation: Wald statistic=1.21, both df=9 and p>0.9).

### Phlyogenetic and statistical analyses

In order to account for the non-independence of data from related species, we first formed groups within which adhesive pads are likely homologous, based on their position on the leg and their structure (i. e. hairy vs. smooth). These groups are (1) Squamata, (2) Anura, (3) Araneae, (4) mites with smooth pads, (5) mites with hairy pads, (6) insects with tarsal hair fields (e. g. some Coleoptera and Raphidioptera), (6) insects with smooth pulvilli (e. g. some Hemiptera), (7) insects with hairy pulvilli (e. g. some Diptera), (8) insects with unfoldable arolia (some Hymenoptera), (8) insects with non-eversible arolia (e. g. some Polyneoptera, Hemiptera and Lepidoptera), (9) insects with specialised distal euplantula (some Polyneoptera), (10) insects with tibial pads (e. g. some aphids). These groups were all connected directly to the root of the tree, so that analogous structures share no branch, and thus the respective elements in the error covariance matrix of our linear models are zero [108]. Within this groups, we assembled a tree topology from phylogenies published for the constituent groups (Anura [109], Squamata [110], Aranae [28], Insecta [111], Blattodea [112], Coleoptera [113], Diptera [114], Hemiptera [115, 116], Hymenoptera [117, 118]. As statistically supported branch lengths were not available, comparative phylogenetic procedures which allow for more complex evolutionary models, such as Ornstein-Uhlenbeck models [119, 120], were not feasible. Instead, we performed phylogenetic generalised least squares on 10.000 trees with randomised branch lengths that were rendered ultrametric via a correlated rate model [121]. In order to account for the uncertainty of the phylogenetic error covariance structure, Pagel’s λ was estimated simultaneously via a maximum likelihood optimisation [17,122]. The fitted coefficients were normally distributed, with a coefficient of variation below 1% (see fig. S1 in Supplementary Information). For simplicity, we report results for an ultrametric tree calculated from a tree with all branch lengths set to 1.

All analyses involving pad area and weight were performed on log_10_-transformed values, and with the the routines implemented in the R packages nlme v.3.1-118, ape v.3.1-1, and phytools v.0.3-93 in R v.3.1.3 [123,124]. There is some controversy as to whether (reduced) major axis or ordinary least squares regression is more appropriate for estimating allometric coefficients from data containing both ‘measurement’ and ‘biological’ error [18, 19, 125, 126, 127]. We thus report results for both techniques, and note that the key results of this study hold independent of what regression model is applied.

An alternative method to account for relatedness is to monitor the change of the estimated parameters as one moves up in taxonomic rank. We performed multiple reduced major axis regressions across all taxa, and separately within the taxonomic levels class, order, family and genus (for example, all Hymenoptera provide one allometric slope data point within the level ‘order’). Within the taxonomic levels, groups were only included if the weight range of the available species exceeded a factor of 3, and if data for at least 4 different groups from the next sub-level were available (e. g. an order was included if at least four families were represented).

In order to visualise the effect of evolutionary history on body size-corrected pad area, residuals from a phylogenetic reduced major axis regression were used to estimate maximum likelihood ancestral states for all nodes and along the branches via the method described in [128]. These values are only rough estimates, first because we do not have statistically supported branch lengths, second because the species sampling in our phylogeny is incomplete, third because we do not account for the influence of relative pad area on diversification rate, and fourth, because only shared ancestors among lower taxonomic ranks possessed adhesive pads. Ancestral state estimates are solely used to visualise systematic differences at the tip-level, and no conclusions are based on them.

### Scaling of pad performance in tree frogs

All data are from whole-animal force measurements, conducted using a tilting platform. In total, we extracted data for pad area and body weight for 17 species belonging to 4 families from references [35,36,37,38]. The phylogenetic tree underlying the phylogenetic regressions was extracted from the detailed phylogeny in [109], and is shown in fig. S2 in the Supplementary Information.

## Acknowledgments

We are sincerely grateful to all our colleagues who readily shared published and unpublished data with us: Aaron M. Bauer, Jon Barnes, Niall Crawford, Thomas Endlein, Hanns Hagen Goetzke, Thomas E. Macrini, Anthony P. Russell & Joanna M. Smith. This study was supported by research grants from the UK Biotechnology and Biological Sciences Research Council (BB/I008667/1) to WF, the Human Frontier Science Programme (RGP0034/2012) to DI, AJC and WF, the Denman Baynes Senior Research Fellowship to DL and a Discovery Early Career Research Fellowship (DE120101503) to CJC.

## Appendix

The following species were included:

**Amphibia**: Anura, Dendrobatidae: Mannophryne trinitatis (Garman, 1887), Hemiphractidae: Flectonotus fitzgeraldi (Parker, 1933), Hylidae: Dendropsophus microcephalus (Cope, 1886); Dendropsophus minusculus (Rivero, 1971); Dendropsophus minutus (Peters, 1872); Hyla cinerea (Schneider, 1799); Hyla versicolor LeConte, 1825; Hypsiboas boans (Linnaeus, 1758); Hypsiboas crepitans (Wied-Neuwied, 1824); Hypsiboas geographicus (Spix, 1824); Hypsiboas punctatus (Schneider, 1799); Litoria caerulea (White, 1790); Osteopilus septentrionalis (Dum´eril & Bibron, 1841); Phyllodytes auratus (Boulenger, 1917); Phyllomedusa trinitatis Mertens, 1926; Scinax ruber (Laurenti, 1768); Smilisca phaeota (Cope, 1862); Sphaenorhynchus lacteus (Daudin, 1801); Trachycephalus venulosus (Laurenti, 1768), Craugastoridae: Pristimantis euphronides (Schwartz, 1967), Ranidae: Staurois gut-tatus (Gu¨nther, 1858), Rhacophoridae: Rhacophorus pardalis Gu¨hnter, 1858

**Arachnida:** Araneae, Ctenidae: Ctenus curvipes (Keyserling, 1881); Ctenus sinuatipes Pickard-Cambridge, 1897; Ctenus sp. 3 Walckenaer, 1805; Cupiennius coccineus Pickard-Cambridge, 1901; Cupiennius getazi Simon, 1891; Cupiennius salei (Keyserling, 1877); Phoneutria boliviensis (Pickard-Cambridge, 1897), Philodromidae: Philodromus aureolus (Clerck, 1757); Philodromus dispar Walckenaer, 1826, Salticidae: Evarcha arcuata (Clerck, 1757); Marpissa muscosa (Clerck, 1757); Salticus scenicus (Clerck, 1757); Sitticus pubescens (Fabricius, 1775); Pseudeuophrys lanigera (Simon, 1871), Theraphosidae: Aphonopelma seemanni (Pickard-Cambridge, 1897), Thomisidae: Misumenops spec. Mesostigmata, Laelapidae: Androlaelaps schaeferi (Till, 1969)

**Trombidiformes, Tetranychidae:** Tetranychus cinnabarinus Boudreaux, 1956; Tetranychus urticae (Koch, 1836)

**Insecta:** Blattodea, Blaberidae: Gromphadorhina portentosa (Schaum, 1853); Nauphoeta cinerea (Olivier, 1789), Blattellidae: Blattella germanica Linnaeus, 1767, Blattidae: Blatta orientalis Linnaeus, 1758; Periplaneta americana (Linnaeus, 1758); Periplaneta australasiae Fabricius, 1775; Supella supellectilium (Serville, 1839)

**Coleoptera, Brentidae:** Cylas puncticollis (Boheman, 1833), Cantharidae: Cantharis rustica Fall´en, 1807; Rhagonycha fulva Scopoli, 1763, Cerambycidae: Agrianome Spinicollis (Macleay, 1827); Clytus arietis (Linnaeus, 1758), Chrysomel-idae: Altica lythri Aub´e, 1843; Cassida canaliculata Laicharting, 1781; Chrysolina americana (Linnaeus, 1758); Chrysolina fastuosa Scopoli, 1763; Chrysolina menthastri (Suffrian, 1851); Chrysolina polita (Linnaeus, 1758); Clytra quadripunctata Linnaeus, 1758; Cryptocephalus spec. Geoffroy, 1762; Galerucella nymphaeaei (Linnaeus, 1758); Gastrophysa viridula (De Geer, 1775); Hemisphaerota cyanea (Say, 1824); Leptinotarsa decemlineata Say, 1824; Oulema melanopus (Linnaeus, 1758); Psylliodes chrysocephalus (Linnaeus, 1758), Cleridae: Trichodes alvearius (Fabricius, 1792), Coccinellidae: Adalia bipunctata (Linnaeus, 1758); Coccinella septempunctata (Linnaeus, 1758); Harmonia axyridis (Pallas, 1773); Henosepilachna vigintioctopunctata (Fabricius, 1775); Psyllobora vigintiduopunctata (Linnaeus, 1758); Subcoccinella vigintiquatuorpunctata (Linnaeus, 1758), Pyrochroidae: Pyrochroa coccinea Linnaeus, 1762, Silphidae: Nicrophorus spec. Fabricius, 1775

**Dermaptera, Forficulidae:** Forficula auricularia Linnaeus, 1758

**Diptera, Calliphoridae:** Calliphora vicina Robineau-Desvoidy, 1830; Calliphora vomitoria (Linnaeus, 1758); Lucilia caesar (Linnaeus, 1758), Syrphidae: Episyrphus balteatus (De Geer, 1776); Eristalis tenax (Linnaeus, 1758); Myathropa florea (Linnaeus, 1758); Sphaerophoria scripta (Linnaeus, 1758); Syrphus ribesii (Linnaeus, 1758); Volucella pellucens (Linnaeus, 1758), Tabanidae: Tabanus spec Linnaeus, 1758, Tachinidae: Tachina fera (Linnaeus, 1761)

**Hemiptera, Aphididae:** Aphis fabae Scopoli, 1763; Megoura viciae Buckton, 1876, Cicadellidae: Aphrodes sp. Curtis, 1833; Eupteryx aurata (Linnaeus, 1758), Coreidae: Coreus marginatus (Linnaeus, 1758); Gonocerus acuteangulatus (Goeze, 1778); Leptoglossus occidentalis Heidemann, 1910, Delphacidae: Asiraca clavicornis (Fabricius, 1775); Javesella pellucida (Fabricius, 1794); Ribautodelphax spec (Ribaut, 1953); Stenocranus minutus (Fabricius, 1787), Heterogastridae: Heterogaster urticae (Fabricius, 1775), Miridae: Dicyphus errans (Wolff, 1804); Lygocoris pabulinus (Linnaeus, 1761), Pentatomidae: Aelia acuminata (Linnaeus, 1758); Palomena prasina (Linnaeus, 1761), Triozidae: Trioza urticae (Linnaeus, 1758) Hymenoptera, Formicidae: Atta cephalotes (Linnaeus, 1758); Atta colombica (Gu´erin-M´eneville, 1844); Camponotus schmitzi St¨arcke, 1933; Myrmica scabrinodis Nylander, 1846; Oecophylla smaragdina Fabricius, 1775; Polyrhachis dives Smith, 1857, Vespidae: Vespa crabro Linnaeus, 1758

**Lepidoptera, Tortricidae:** Cydia pomonella (Linnaeus, 1758) Mantodea, Mantidae: Stagmomantis theophila Rehn, 1904 Orthoptera, Acrididae: Chorthippus brunneus (Thunberg, 1815); Locusta migratoria manilensis (Linnaeus, 1758), Tettigoniidae: Conocephalus discolor (Thunberg, 1815); Metrioptera roeselii (Hagenbach, 1822)

**Phasmatodea, Bacillidae:** Heteropteryx dilatata (Parkinson, 1798), Diapheromeridae: Carausius morosus Sinety, 1901; Clonaria conformans (Brunner, 1907), Heteropterygidae: Trachyaretaon carmelae Lit & Eusebio, 2005, Phasmatidae: Eurycantha calcarata (Lucas, 1869); Medauroidea extradentata Brunner, 1907; Ramulus spec Saussure, 1862; Sceptrophasma hispidulum Wood-Mason, 1873 Raphidioptera, Raphidiidae: Phaeostigma notatum (Fabricius, 1781)

**Mammalia:** Chiroptera, Myzopodidae: Myzopoda aurita Milne-Edwards & A. Grandidier, 1878

**Reptilia:** Squamata, Dactyloidae: Anolis auratus Daudin, 1802; Anolis biporcatus Wiegmann, 1834; Anolis capito (Peters, 1863); Anolis carolinensis Voigt, 1832; Anolis cristatellus (Dum´eril & Bibron, 1837); Anolis cupreus (Hallowell, 1860); Anolis distichus (Cope, 1861); Anolis equestris (Merrem, 1820); Anolis evermanni (Stejneger, 1904); Anolis frenatus (Cope, 1899); Anolis garmani (Stejneger, 1899); Anolis grahami (Gray, 1845); Anolis humilis (Peters, 1863); Anolis leachi (Dum´eril & Bibron, 1837); Anolis lemurinus (Cope, 1861); Anolis limifrons (Cope, 1871); Anolis lineatopus Gray, 1840; Anolis li-onotus (Cope, 1861); Anolis pentaprion (Cope, 1863); Anolis poecilopus (Cope, 1862); Anolis polylepis (Peters, 1874); Ano-lis pulchellus Dum´eril & Bibron, 1837; Anolis sagrei (Dum´eril & Bibron, 1837); Anolis valencienni (Dum´eril & Bibron, 1837), Diplodactylidae: Correlophus ciliatus Guichenot, 1866, Gekkonidae: Chondrodactylus bibronii (Smith, 1846); Gehyra mutilata (Wiegmann, 1834); Gehyra oceanica (Lesson 1830); Gehyra vorax Girard, 1858; Gekko athymus Brown & Alcala, 1962; Gekko gecko (Linnaeus, 1758); Gekko gigante Brown & Alcala, 1978; Gekko japonicus (Schlegel, 1836); Gekko mindorensis Taylor, 1919; Gekko monarchus (Taylor, 1917); Gekko palawanensis Taylor, 1925; Gekko romblon (Brown & Alcala, 1978); Gekko Smithii Gray, 1842; Gekko subpalmatus (Gu¨nther, 1864); Gekko swinhonis Gu¨nther 1864; Gekko vittatus Houttuyn, 1782; Hemidactylus frenatus Schlegel, 1836; Hemidactylus turcicus (Linnaeus, 1758); Lepidodactylus lugubris (Dum´eril & Bibron, 1836); Lepidodactylus pumilus (Boulenger, 1885); Phelsuma dubia (Boettger, 1881); Phelsuma grandis Gray, 1870; Phelsuma laticauda (Boettger, 1880); Rhoptropus afer Peters, 1869; Rhoptropus barnardi Hewitt, 1926; Rhoptropus biporosus FitzSimons, 1957; Rhoptropus boultoni Schmidt, 1933; Rhoptropus bradfieldi Hewitt, 1935; Rhoptropus cf biporosus (Fitzsimons,1957); Rhoptropus diporus Haacke, 1965, Scincidae: Prasinohaema virens Peters, 1881; Prasinohaema prehensicauda (Loveridge, 1945); Prasinohaema flavipes (Parker, 1936); Lipinia leptosoma (Brown & Fehlmann, 1958)

